# A simple neural circuit model explains diverse types of integration kernels in perceptual decision-making

**DOI:** 10.1101/2024.12.10.627688

**Authors:** Xuewen Shen, Fangting Li, Bin Min

## Abstract

The ability to accumulate evidence over time for deliberate decision is essential for both humans and animals. Decades of decision-making research have documented various types of integration kernels that characterize how evidence is temporally weighted. While numerous normative models have been proposed to explain these kernels, there remains a gap in circuit models that account for the complexity and heterogeneity of single neuron activities. In this study, we sought to address this gap by using low-rank neural network modeling in the context of a perceptual decision-making task. Firstly, we demonstrated that even a simple rank-one neural network model yields diverse types of integration kernels observed in human data—including primacy, recency, and non-monotonic kernels—with a performance comparable to state-of-the-art normative models such as the drift diffusion model and the divisive normalization model. Moreover, going beyond the previous normative models, this model enabled us to gain insights at two levels. At the collective level, we derived a novel explicit mechanistic expression that explains how these kernels emerge from a neural circuit. At the single neuron level, this model exhibited heterogenous single neuron response kernels, resembling the diversity observed in neurophysiological recordings. In sum, we present a simple rank-one neural circuit that reproduces diverse types of integration kernels at the collective level while simultaneously capturing complexity of single neuron responses observed experimentally.

**Author Summary:** This study introduces a simple rank-one neural network model that replicates diverse integration kernels—such as primacy and recency—observed in human decision-making tasks. The model performs comparably to normative models like the drift diffusion model but offers novel insights by linking neural circuit dynamics to these kernels. Additionally, it captures the heterogeneity of single neuron responses, resembling diversity observed in experimental data. This work bridges the gap between decision-making models and the complexity of neural activity, offering a new perspective on how evidence is integrated in the brain.

## Introduction

Accumulating evidence over time is crucial for deliberate decision-making. The underlying neural mechanisms have been intensively investigated in the literature [1–9]. Classic decision-making theory predicts equal weighting of sensory evidence over time [6, 10]. However, humans and animals frequently apply uneven temporal weighting to sensory evidence. For instance, previous studies have shown that early evidence exerts larger effect than the latter one on the final decision, a phenomenon referred to as primacy effect [11–12]. Conversely, other studies have found evidence supporting the recency effect, where later evidence has a larger impact [13–14]. More dramatically, recent findings suggest that non-monotonic temporal weighting profiles may also be prevalent under certain conditions [15–17]. To explain this diversity of integration kernels, numerous models, including the drift-diffusion model [15, 18–20] and the divisive normalization model [17], have been proposed. Despite the successful explanation of the behavioral data, recorded single-neuron responses in experiments oftentimes exhibit significant heterogeneity [21–23]. The underlying neural mechanisms and their relationship with behavior require further investigation.

In the present study, we sought to investigate the neural mechanisms underlying the diversity of integration kernels in perceptual decision-making using a low-rank recurrent neural network (RNN) modeling approach. Low-rank RNN is a fruitful modeling framework not only accounting for single neuron response complexity but also retaining the collective-level mechanistic transparency [24–25]. This approach has proven incredibly valuable in modeling various cognitive process, including perceptual decision-making, parametric working memory [25], context-dependent decision-making [26] and timing [27]. By leveraging the expressive and interpretable power of low-rank RNNs, we developed a unified yet minimal model to account for various integration kernels in perceptual decision-making. Furthermore, we strived to go beyond the conceptual understanding and tested the capacity of low-rank RNNs in fitting the behavioral data in experiments. When applied to the human behavioral data [16], even the simplest rank-one model demonstrated fitting performance comparable to state-of-the-art models, including the drift-diffusion and divisive normalization models. Furthermore, going beyond the previous normative models (e.g., [17]), single neurons in our model exhibited heterogeneous response kernels, resembling the complex single neuron responses observed in experiments. In summary, we presented a simple neural circuit model with rank-one dynamics that not only reproduces diverse types of integration kernels and quantitatively accounts for the human behavioral data, but also accommodates the complexity of single neuron responses observed in neurophysiological experiments.

## Results

### The task paradigm and the modeling approach

The task we studied in this work is the click-version perceptual decision-making task (Fig 1a, [16–17]). In this task, participants were presented with a series of 20 clicks delivered over 1 second at a frequency of 20 Hz during each trial. The clicks were presented on either the left or right side. After the sequence, participants had to determine which side had more clicks. Using this task paradigm, Keung et al. [17] identified four “behavioral phenotypes”, each exhibiting a distinct integration kernel profile, including flat, recency, primacy and bump ones (Fig 1b and S1 Fig).

**Fig 1.**
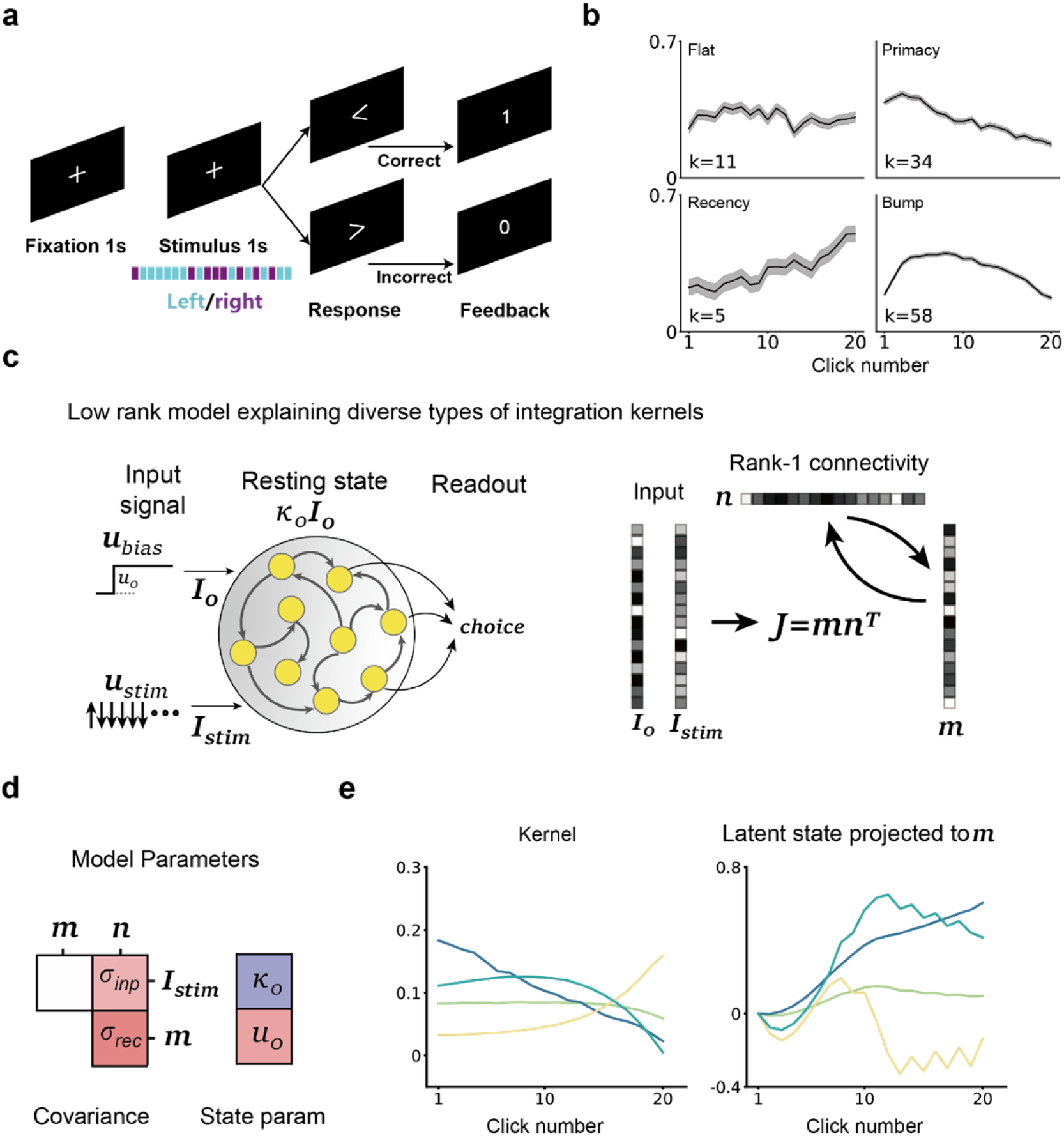
A rank-1 network model accounts for diverse types of integration kernels in click-version perceptual decision task. **a.** Click-version perceptual decision-making task paradigm (adapted from [17]). Human participants listen to a train of 20 clicks for 1 second. Each click is presented either on left side (cyan colored) or right side (purple colored). At the end of trial, participants report which side had **b.** Human participants’ integration kernels are classified into four types. We categorized 108 participants into four distinct groups based on the shape of their integration kernels, using polynomial fitting (see Supplementary Note 1). The solid black lines represent typical integration kernels for each group of participants, while the shaded areas indicate s.e.m. across participants. **c.** A rank-1 network model performing click-version perceptual decision-making task. Left: The rank-1 network receives two input signals *u*_*bias*_ and *u*_*stim*_. *u*_*bias*_is a step function representing the overall excitation from incoming stimulus with a step size of *u*_0_, while *u*_*stim*_encodes the click input sequence (+ 1 for left click and ― 1 for right click); Right: The connectivity of rank-1 network is characterized by four column vectors: the overall excitation input embedding *I*_0_, the stimulus input embedding *I*_*stim*_, and the recurrent column vectors *n* and *m*. The recurrent connectivity matrix is given by *J* = *mn*^*T*^. **d.** The rank-1 network model includes four free parameters: the input strength σ_*inp*_ defined as the covariance between *n* and *I*_*stim*_; the recurrent strength σ_*rec*_ defined as the covariance between *n* and *m*; the initial state parameter *κ*_0_and the overall excitation parameter *u*_0_. **e.** The rank-1 networks can reproduce various types of integration kernels. Left: Different models simulate flat, primacy, recency, bump integration kernels; Right: The latent state projected ont *m*, representing the accumulation of incoming auditory evidence (positive for left side evidence and negative for right side evidence) over the course of a trial, as shown in Fig 1a.

To investigate the neural mechanisms underlying these “behavioral phenotypes”, we modeled the same task using the low-rank RNN modeling approach [24–25]. Specifically, we constructed a rank-1 neural network model with N neurons (Fig 1c, Left, N = 512 in this study) described by the following equation

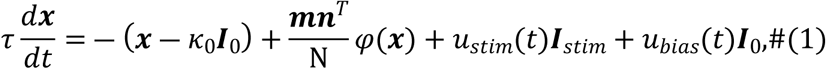

where *x* is a N-dimensional vector representing the hidden state of the network, τ is the single neuron time constant and φ is the nonlinear activation function (*tanh* in this article recurrent connectivity, which can be written as the outer product of two N-dimensional column vectors *m* and *n* (Fig 1c, right).

Before the stimulus presentation, the network remains at the initial state along the direction *I*_0_with amplitude *κ*_0_. During the stimulus period, a pulse input *u*_*stim*_(*t*) mimicking the auditory pulse sequence, and a bias input with amplitude *u*_0_ mimicking the overall excitation, are applied to the network via the embedding vectors *I*_*stim*_and *I*_0_, respectively. Note that the connectivity vectors in this model, including *m*, *n*, *I*_*stim*_, *I*_0_, were generated according to a zero-mean Gaussian distribution as typically assumed in low-rank neural network models [24–25] (see Methods for details). After the auditory sequence is presented, the decision probability is then formed based on the decision variable *κ*_*dv*_(*t*) = *x*(*t*) ⋅ *m*/N at the choice time *T*. That is, *logit*(*p*_*left*_ at trial *k*) = *K*_*dv*_ (*T*).

### Understanding the decision variable dynamics at the collective level

Endowed by the mean-field theory of low-rank neural networks [24–25], when the neuron number N becomes sufficiently large, the dynamics of decision variable can be approximated by the following equation

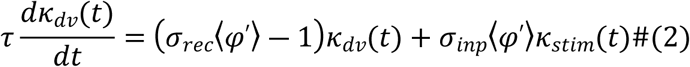

where σ_*rec*_denotes the overlap between connectivity vectors *m* and *n*, σ_*inp*_represents the overlap between connectivity vectors *I*_*stim*_and *n*. Here, *κ*_*stim*_(*t*) is the low-pass filtered version of the pulse input *u*_*stim*_(*t*) (see Eq. 6 a in Methods), and ⟨φ′⟩ is the average gain of single neuron in the network. Unlike previous models (e.g., the divisive normalization model in [17]) that focus only on gain modulation of the input (the second term on the right-hand side), this model shows that gain modulation also impacts the effective recurrent strength (the first term on the right-hand side). Importantly, the average gain ⟨φ′⟩ depends not only on the decision variable *κ*_*dv*_(*t*) along the axis *m* per se but also on the activities along all other axes, including *κ*_*stim*_(*t*) along the axis *I*_*stim*_and the activity (denoted as *κ*_*bias*_(*t*)) along the axis *I*_0_(see Methods for more details). Therefore, while the activity *κ*_*bias*_in the axis *I*_0_does not directly drive the evolution of the decision variable, it can modulate the integration process and thereby the integration kernel.

Guided by Eq. 2, by varying four parameters—σ_*rec*_, σ_*inp*_, *κ*_0_ and *u*_0_ (Fig 1d)—we can replicate all four types of integration kernels (Fig 1e) observed in human behavioral data.

### An explicit expression of integration kernels

The collective dynamics described by Eq. 2 not only allow for an exploration of the rank-1 neural network’s capability in generating various integration kernels but also provide an explicit expression for these kernels. Specifically, the formal solution for decision variable *K* as function of time *T* can be written as: 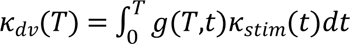, where

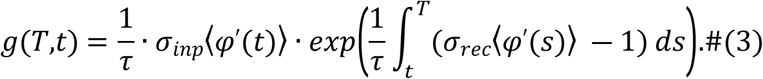

That is, the integration kernel *g*(*T,t*) is the product of the effective instantaneous input strength σ⟨φ′(*t*)⟩ and the effective temporal factor 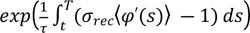 scaled by time constant τ (see Methods for the derivation). This expression provides a clear and intuitive understanding of how each click at different time points contributes to the decision.

Using this expression, we investigated the mechanisms underlying the generation of different integration kernels in our model. For the flat kernel, as expected, both the effective instantaneous input strength and the effective temporal factor exhibited minimal variation over time, the combination of which led to a flat profile (Fig 2a, top). For the primacy (recency) kernel, typically, both the effective instantaneous input strength and the effective temporal factor were decreasing (increasing) over time (see additional varieties in S2 Fig and Supplementary Note 2), producing a decreasing (increasing) profile over time (Fig 2a, middle and bottom). Regarding the bump kernel, we observed that a combination of a decreasing effective instantaneous input strength profile and an increasing effective temporal factor profile can produce such a kernel type (Fig 2b, top). This mechanism is consistent with the divisive normalization model [17] where the input strength was down-regulated by a gain control unit over time, while an increasing temporal factor is assumed to be independent of gain modulation. However, in our low-rank model family, such a combination is not the only possibility leading to a bump kernel. For instance, an increasing effective input strength combined with a bump-shaped effective temporal factor can also yield a bump kernel (Fig 2b, bottom). How is it possible having a bump-shaped effective temporal factor in our low-rank models? The explicit expression of the effective temporal factor 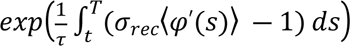 provides us a concise explanation. Let us consider the scenario of increasing temporal gain. In this scenario, the term σ_*rec*_⟨φ^′^(*S*)⟩ ― 1 is increasing over time. Therefore, it is possible to have σ_*rec*_⟨φ^′^(*S*)⟩ ― 1 less than 0 at the initial time, cross 0 at certain time point during the trial and eventually greater than 0 by the final choice time. As a consequence, the integral 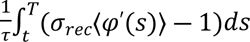 as a function of time *t* will first increase, achieve the peak value at the time when σ_*rec*_⟨φ^′^(*S*)⟩ ― 1 is equal to 0, and then decease, thereby giving rise to a bump profile of the effective temporal factor. This kind of mechanism essentially relies on the transition from a leaky integrator (i.e., σ_*rec*_⟨φ^′^(*S*)⟩ ― 1 < 0) to an unstable integrator (i.e., σ_*rec*_⟨φ^′^(*S*)⟩ ― 1 > 0) during the trial. We verified that this mechanism of bump kernel also applies to low-rank models with different activation functions (S2b Fig and Supplementary Note 3), demonstrating the generality of this mechanism.

**Fig 2.**
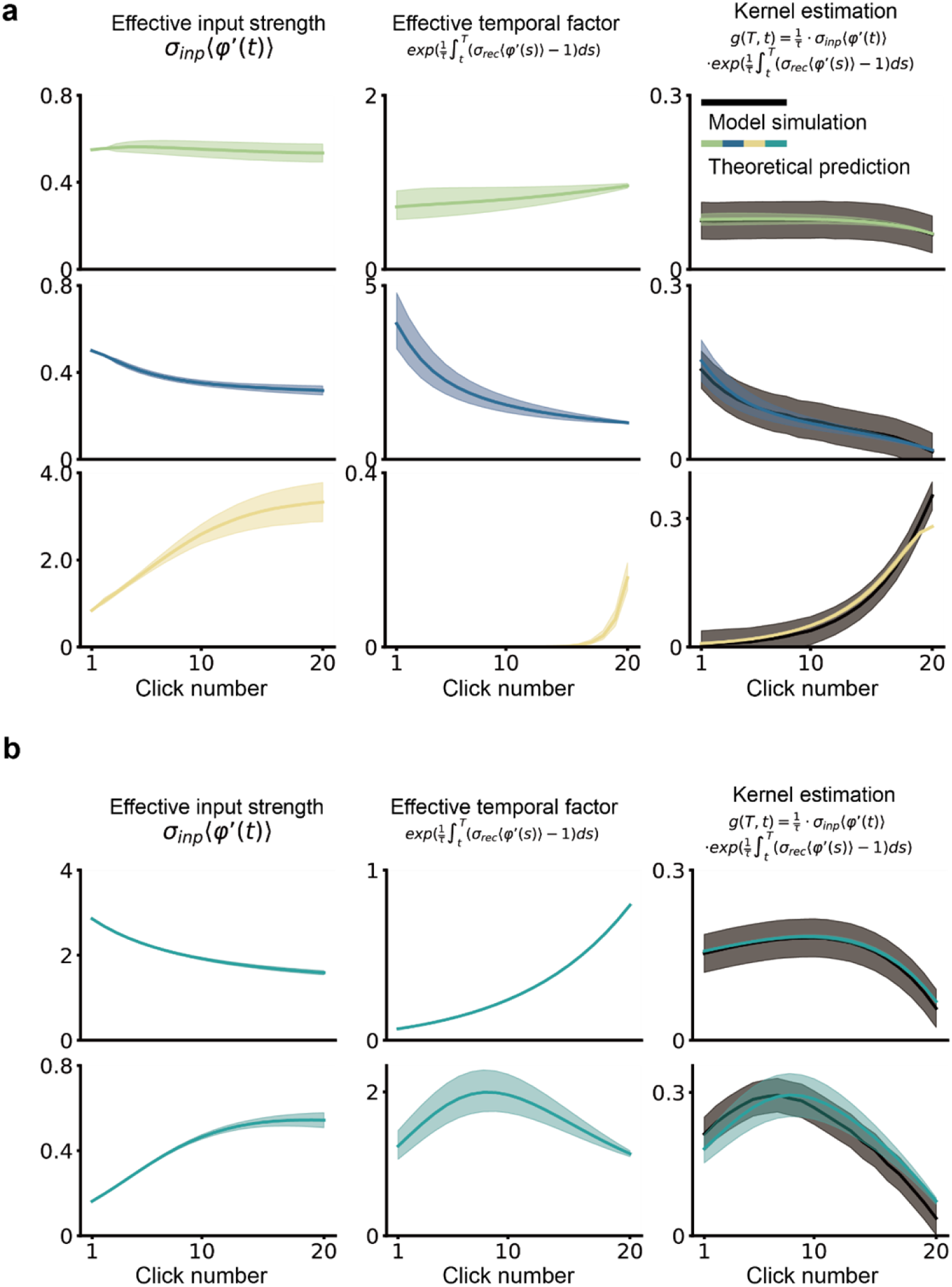
Collective dynamics reveal mechanisms underlying different kernel shapes. **a.** Example model simulations (first row, flat; second row, primacy; third row, recency) illustrate how integration kernels (right column) arise from the multiplicative interactions between two components—the effective input strength (first column) and the effective temporal factor (middle column). In the right column, black lines represent the integration kernels from model simulations while colored lines indicate the theoretical prediction (i.e., Eq. 3). The integration kernel is obtained through regression analysis for the behavioral outputs generated from model simulations (see Eq. 1a in Methods for more details). The shaded areas denote s.e.m. across trials. **b.** Simulations of two distinct models demonstrate different implementations of bump-shaped kernels with rank-1 network models. First row: A monotonically decreasing effective input strength combined with an increasing effective temporal factor results in a bump kernel. Second row: A monotonically increasing effective input strength combined with a bump-shaped effective temporal factor also generates a bump kernel.

Notably, the estimation of integration kernel does not depend on how we readout choices from rank-1 network model, whether from hidden states or neural firing rates. We simulate choices with linear readout of neural firing rates in S3 Fig following the decision probability: *logit*(*p*_*left*_ at trial *k*) = φ (*x*(*t*)) ⋅ *W*/N at choice time T, where *W* represents a N-dim readout vector.

In summary, the collective-level analysis of low-rank models reveals a concise mechanism unifying multiple types of integration kernels and helped identify a novel mechanism deviating from the divisive normalization explanation for the intriguing non-monotonic bump kernel.

### Rank-1 network models can quantitatively account for the human behavioral data

Having demonstrated that the rank-1 network models can qualitatively reproduce diverse types of integration kernels, we then asked how well this kind of network models can account for the human behavioral data (i.e., the data presented in [16]). To this end, we followed the same data fitting procedure used in the divisive normalization model [17]. Specifically, in our model, we assumed that a choice is made by comparing the decision variable to noise (parameterized by σ_0_), with an additional decision bias β. As did in [17], we also include an offset parameter μ in the integration kernel. Thus, the probability of choosing left at trial *k* occurring to data fitting process is given by

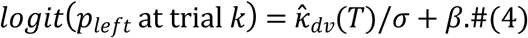

where 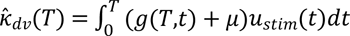. This approach allows us to quantitatively compare model’s predictions with human behavioral data.

The model comprises seven parameters in total: the recurrent strength σ_*rec*_, the input strength σ_*inp*_, the initial state parameter *κ*_0_, the overall excitation during input presentation *u*_0_, the noise level σ, the offset μ and the decision bias β. We fitted the model to human participant data from [16] by maximizing the log-likelihood function (see Methods for details). Our results indicate that rank-1 network models can quantitatively account for all types of integration kernels observed in human data (Fig 3a). By applying the optimal parameters derived from fitting individual participants’ data,

**Fig 3.**
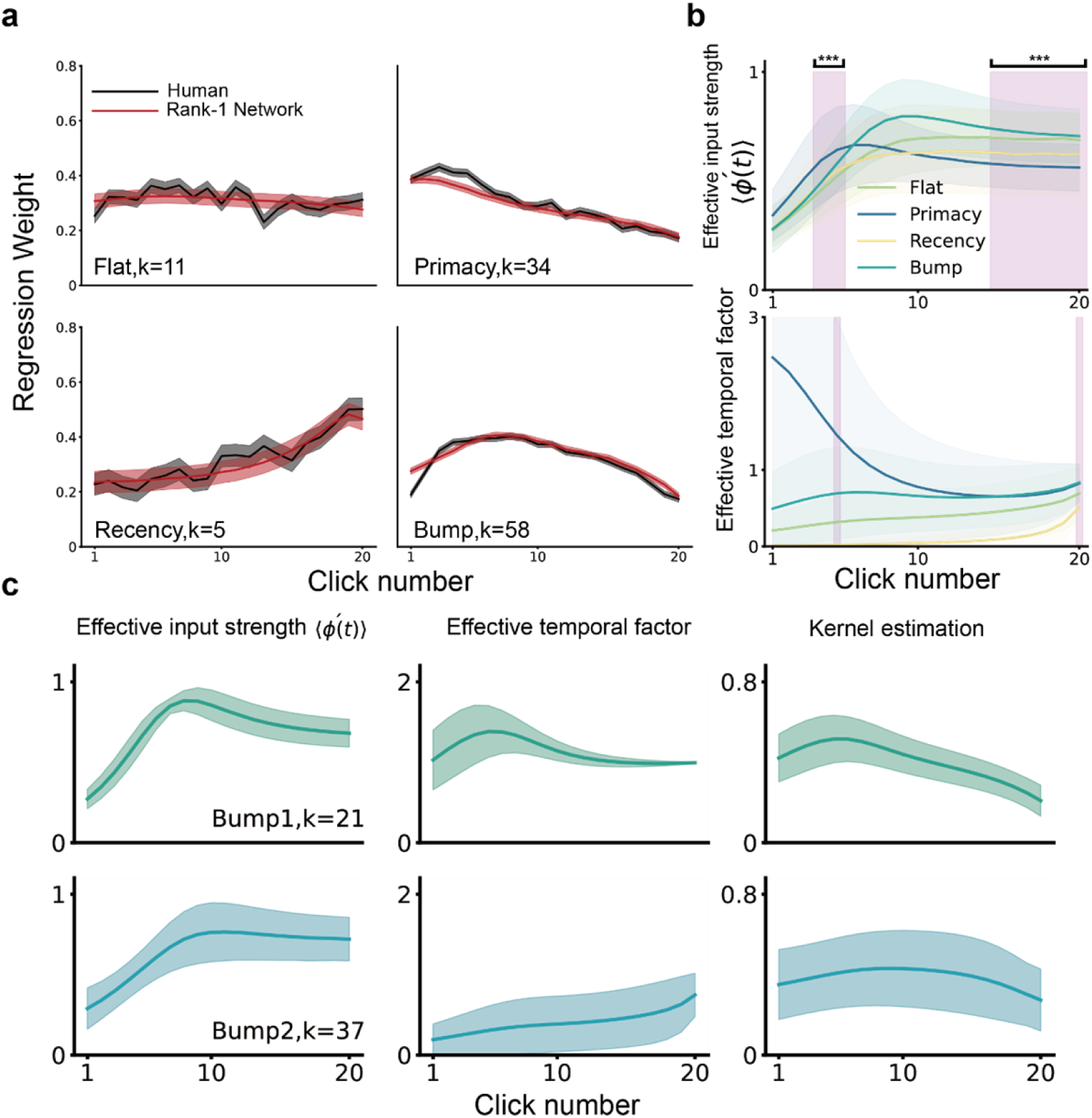
Rank-1 network model accounts for distinct human integration kernels. **a.** Rank-1 networks optimized to fit human participants’ choices. Black lines indicate human integration kernels. Red lines indicate integration kernels generated by rank-1 network models. All shaded areas indicate s.e.m. across participants grouped in the same kernel types. **b.** Optimized model simulation of participant effective input strength and effective temporal factor varies across groups. **Top**: Effective input strength ⟨φ^′^(*t*)⟩ across participants grouped in the same kernel types. The difference of input modulation between groups are significant (***: multivariate ANOVA, Wilk’s lambda, F(60, 254)=2.17,p<0.001). Post-hoc Tukey’s test shows that difference arises from significant higher ⟨φ^′^(*t*)⟩ of primacy kernel compared with flat and bump kernel at 4th and 5th click, and on the contrary, significant lower ⟨φ^′^(*t*)⟩ of primacy kernel compared with flat and bump kernel from 15th to 20th click. The significant area is marked as magenta shaded area. **Bottom**: Effective temporal factor across participants grouped in the same kernel types. Primacy and recency kernels exhibit distinct and apparent trends of decaying and rising temporal weights in cumulative effect. Flat kernel is relatively smooth and balanced between primacy and recency kernel. Bump kernel has two peaks at 5th and 20th click which are marked as magenta shaded areas. **c.** Detailed Categorization of bump shaped kernels. The first row indicates integration of bump shaped kernels with peak of effective temporal factor before 10-th click (21 participants). The second row indicates integration of bump shaped kernels with later peaks (37 participants). The three columns indicate effective input strength, effective temporal factor and estimated kernels respectively. The colored lines are group-averaged and the shaded areas are s.e.m..

we reconstructed the rank-1 network model for each participant. We then simulated choices to generate integration kernels and psychometric curves (S4 and S5 Fig), which closely matched the human data. The distributions of the best-fitted network model parameters are shown in S6 Fig.

These fitted network models also provided us an opportunity to better investigate the potential underlying mechanisms for the different integration kernels observed in the human data. First, we found that the effective input strength generally increases over time (Fig 3b, top), which contrasts with the divisive normalization model, where the effective input strength decreases over time [17]. In addition, we found that the primary distinction between different kernels is reflected in the profile of the effective temporal factor (Fig 3b, bottom). More specifically, for the primacy (recency) kernel, the effective temporal factor exhibited a decreasing (increasing) profile, which is consistent with our intuition. For both flat and bump kernels, on average, the effective temporal factor exhibited minimal variations over time. To further elucidate the mechanisms underlying bump kernels, we categorized the fitted models with bump kernels based on the peak time point of the effective temporal factor (see Supplementary Note 1 for more details). This classification revealed two distinct groups: one with the peak occurring in the middle of the trial (Fig 3c, top) and another with the peak at the end of the trial (Fig 3c, bottom), suggesting that multiple mechanisms may underlie the same integration kernel type, both in theory and in the empirical data.

### Rank-1 network models performs as well as state-of-art models

We then compared the fitting performance between the rank-1 network model with other normative models. The first normative model we considered is the leaky competitive accumulator (LCA) model [28], an extension of the standard drift diffusion model [29]. The LCA model can be written as:

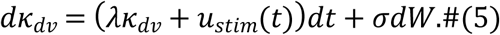

where λ represents the memory drift and σ is the diffusion noise rate of a Wiener process pointed out in [17] this LCA model can account for the flat (λ = 0), recency (λ < 0) and primacy (λ > 0) kernels but is incapable of producing the bump kernel. The second model we compared is the divisive normalization model [17], which can be written as:

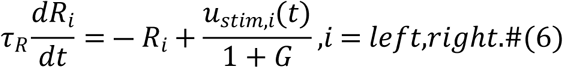

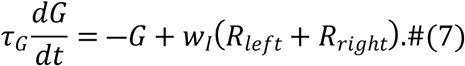

where *R*_*left*_ and *R*_*right*_ represent the accumulated evidence of the left and right side clicks, respectively, *u*_*stim,left*_(*t*) and *u*_*stim*,*right*_(*t*) denote the auditory click input from the left and right sides, respectively, and *G* represents the divisive gain normalizing the click input strength to the accumulator. The choice is determined by comparing *R*_*left*_ ― *R*_*right*_ with a soft bound (see Methods for more details). Previous work demonstrated that this divisive normalization model could account not only for the flat, recency and primacy kernels but also for the bump kernel by varying the parameters τ_*R*_, τ_*G*_ and *W*_*I*_ [17]. By comparing with these two models using log likelihood, Akaike information criterion (AIC) and Bayesian information criterion (BIC), we found that our rank-1 model outperformed the LCA model and performed comparably to the divisive normalization model (Table 1). Simulated integration kernels and psychometric curves for the best-fitted LCA and divisive normalization models are presented in S7, S8, S9 and S10 Fig.

**Table 1.**
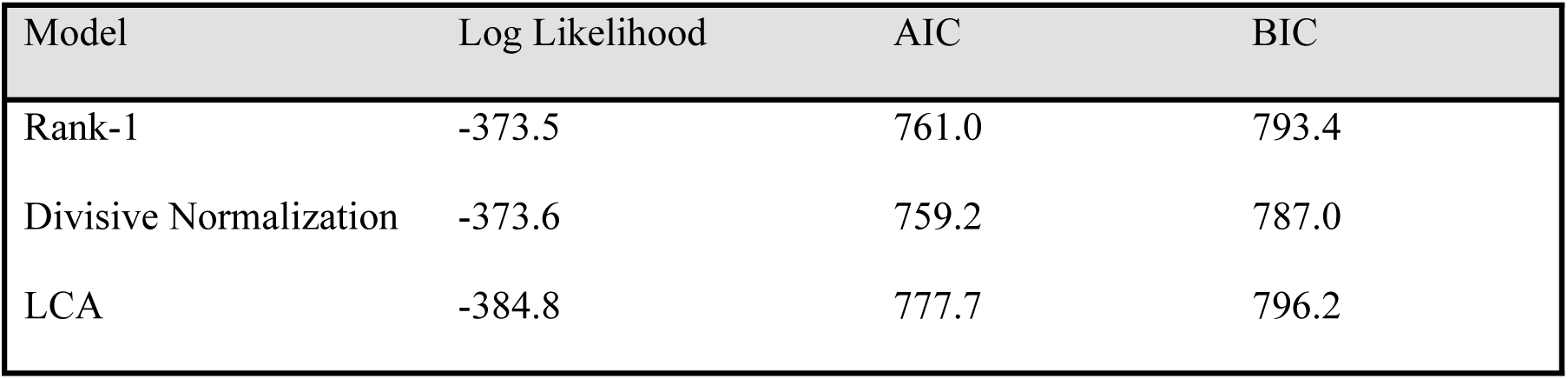
The rank-1 circuit model perform as well as existing models.

| Model | Log Likelihood | AIC | BIC |
| --- | --- | --- | --- |
| Rank-1 | -373.5 | 761.0 | 793.4 |
| Divisive Normalization | -373.6 | 759.2 | 787.0 |
| LCA | -384.8 | 777.7 | 796.2 |

In summary, we showed that the rank-1 model can not only qualitatively generates diverse types of integration kernels in theory but also quantitatively account for real data in a manner comparable with the state-of-the-art normative models.

### The heterogeneity of single neuron integration kernels

Distinct from other normative models, our rank-1 neural network model allowed us to explore the single neuron integration property that can be directly examined in experiments. Similar to previous neurophysiological studies (e.g., [30]), we fitted the firing rate of each single neuron in the network at the choice time as the sum of a stimulus component and a choice component (Fig 4a and S11 Fig). The stimulus component represents the sum of each click stimulus input weighted by corresponding time-dependent regression coefficients. These time-dependent regression coefficients were defined as the single neuron integration kernel, characterizing how clicks at different time points affect the firing rate of a given single neuron at the choice time. Using this definition (see Methods for more details), we computed the integration kernel for each neuron in our model. We found that for any given model in our model family, there was significant variability in integration kernels across neurons. For instance, in a model with a bump kernel (Fig 4b), while some neurons exhibited the typical bump integration kernels, others displayed integration kernels with decreasing or increasing profiles, which were qualitatively different from the bump kernel. What mechanisms underlie this ubiquitous heterogeneity in single neuron responses? The mechanistic transparency of our low-rank models allowed us to investigate this question in detail.

**Fig 4.**
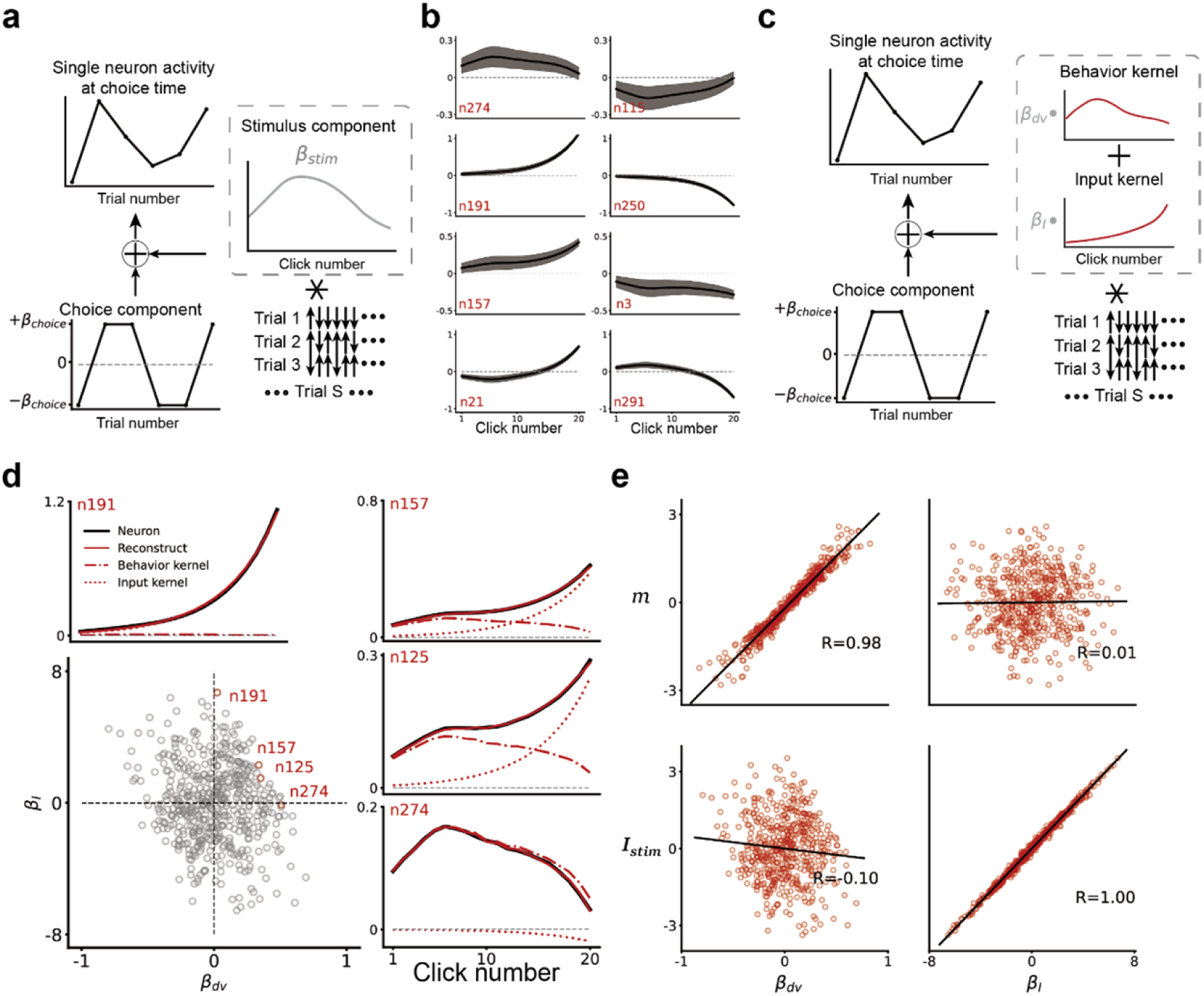
Single neurons in our model exhibited heterogeneous response kernels resembling the diversity of single neuron responses. **a.** Defining single neuron integration kernels through regression. Single neuron activity at choice time is modeled as a combination of a choice component multiplied by the choice of trials and the convolution of a stimulus component (gray dashed-line box) and the click input sequence. The stimulus component, by definition, is the single neuron integration kernel. **b.** Diverse types of single neuron integration kernels. The activity of each neuron in a simulated rank-1 network model which accounts for typical bump kernel (Fig 3a, Bump kernel) at choice time is fitted using choice components and neuron integration kernels. Neurons display diverse types of integration kernels, including behavior kernel stereotypes (n274 and n115 for inverse behavior kernel), input kernel stereotypes (n191 and n250 for inverse input filter) and other shapes (n157, n3, n21 and n291). **c.** Approximating the single neuron integration kernel as the summation of choice component and the convolution of the click input sequence and two components—a behavioral kernel component and an input kernel component (gray dashed-line box). The regression coefficients of neuron *i* for the behavior kernel and input kernel are β^*i*^ and β^*i*^, respectively. **d.** Evaluation of the approximation in (c). The black lines in top and right panels indicate neuron integration kernel by regression model. The red lines indicate the reconstructed single neuron integration kernel, which are linear combinations of the behavior kernel (red dash-dot line) and input kernel (red dotted line). Example neurons are selected from all neurons in the rank-1 network (lower left). Red dots represent selected neurons (n274, n125, n157 and n191), while gray dots represent other neurons in the network. The selected neurons are arranged in descending order based on the relative weights of input kernel (β_*I*_) and behavior kernel (β_*dv*_). **e.** Correlation between network connectivity weights and regression coefficients. We compared *m* and *I*_*stim*_. Each red dot represents a single neuron in rank-1 network. The black lines indicate the regression lines for corresponding pairs of network connectivity and regression coefficient. Significant correlations are observed for two pairs (upper left: *m* and β_*dv*_, k=3.94, R=0.98, p<0.001; lower right: *I*_*stim*_and β_*I*_, k=0.52, R=1.00, p<0.001). The other two pairs show no significant correlation (upper right: *m* and β_*I*_, k=-0.01, R=0.01, p=0.76; lower left: *I*_*stim*_and β_*dv*_, k=-0.49, R=-0.10, p=0.03).

To begin with, recall that in our low-rank models, the hidden state *x* can be decomposed into the concatenation of activities along different axes. More specifically,

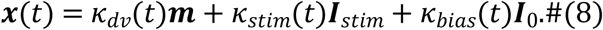

where only *κ*_*dv*_ (determining the behavioral readout) and *κ*_*stim*_ (low-pass filter of the click input) depend on the click input. Therefore, we hypothesized that the single neuron integration kernel, which quantifies the effect of clicks at different time points on the firing rate at the choice time, can be modeled as the sum of a behavioral kernel component and an input kernel component (Fig 4c). Our regression results confirmed this hypothesis, showing that single neuron kernels can indeed be reconstructed by these two components (Fig 4d and S12 Fig, see more details in Methods and Supplementary Note 4). When the single neuron’s firing rate is predominantly influenced by the input kernel component, the integration kernel exhibited an exponential profile. Conversely, when the firing rate is primarily determined by the behavioral kernel component, the integration kernel closely resembled the behavioral kernel profile. For most single neurons, the integration kernels were a mixture of these two components, with the proportion of each component well-predicted by the connectivity strength (Fig 4e). We validated this hypothesis with different activation functions (Supplementary Note 3) and observed similar predictions regarding the connectivity strength (S13 Fig, see more details in Supplementary Note 4).

In summary, our findings demonstrate that the connectivity noise in our low-rank models naturally gives rise to the heterogeneity of single neuron integration kernels. This minimal model resembles the complexity of single neuron responses observed in experiments.

## Discussion

In this work, we introduced a family of simple low-rank neural network models that unify the diverse types of integration kernels observed in human behavioral data. This family of models offers insights at two levels. At the collective level, these models can be regarded as a natural extension of divisive normalization models. They provide a theoretical framework that integrates various types of integration kernels, demonstrating how collective dynamics in neural networks can account for different patterns of evidence accumulation. At the single neuron level, these models reveal a form of single neuron response heterogeneity that mirrors the complex activities observed in experimental studies. Together, our findings suggest that the low-rank neural network modeling framework offers a promising theoretical basis for understanding the underlying circuit mechanisms involved in the versatile process of evidence accumulation over time. By bridging the gap between collective network behavior and individual neuron responses, this approach enhances our ability to elucidate the mechanisms driving decision-making processes.

### Low-rank models as the extension of divisive normalization models

Divisive normalization, as a canonical computation principle [31], has been employed to explain a wide range of cognitive phenomena, ranging from sensory processing in visual systems [32–33] to context-dependent value coding in parietal areas [34]. More recently, it has been applied to perceptual decision-making, successfully explaining the presence of diverse integration kernel types in human data [17]. In this study, we propose that low-rank neural network models can be viewed as a natural extension of divisive normalization models. By conceptualizing divisive normalization as a specific case of gain modulation, our work suggests that these low-rank models provide alternative mechanisms for explaining various integration kernel types, including the bump kernels observed in human data (e.g., Fig 2b, bottom). Future research should focus on designing refined experiments or conducting more detailed analyses to differentiate between these alternative mechanisms and further elucidate the underlying principles.

### The two-level description of the evidence accumulation process over time

In contrast to other normative models [17–18], one distinctive feature of our low-rank models is that it naturally affords a two-level description. Therefore, aside from a similar collective description for the behavioral data to other normative models, our low-rank models provided a closer link with neurophysiological data. For instance, the low-rank framework enables a principled explanation for the observed heterogeneity in single neuron responses within our model (Fig 4). This feature of our model potentially can be extended to explain similar heterogeneous response properties observed in experimental studies.

### Beyond two-alternative perceptual decision-making

While the present study primarily addresses two-alternative perceptual decision-making task, the low-rank modeling framework can be equally applied to multi-alternative decision-making [35–36]. Furthermore, based upon a novel high-throughput animal training paradigm, recent work has identified the fast and slow modes of integration kernels in context-dependent decision-making [30]. The same low-rank modeling framework can be adopted to account for these dynamical modes, offering new insight into the circuit mechanisms underlying the diverse temporal profiles observed in such decision-making processes.

### Limitation of this work

In this study, the low-rank neural network modeling was employed to account for the human behavioral data in a data-driven manner. However, in the current implementation, the neural data has not been incorporated into the model training, thereby not yet fully leveraging the unique advantage of the low-rank neural network modeling as a two-level mechanistic framework. How to incorporate the neural data into the low-rank neural network modeling warrants further systematic investigation [23, 37–39]. Moreover, current model does not account for additional factors contributing to decision such as trial history [40], the distinct roles of multi-regions [39, 41].

In summary, we introduced a neural circuit model not only concisely unifying diverse types of integration kernels but also accommodating the complexity of single neuron responses observed in experiments, making significant contribution towards revealing the neural circuit mechanisms underlying the intriguing evidence accumulation process over time.

## Methods

### Integration Kernel

To characterize each click’s contribution to participants’ choice on each trial, we applied logistic regression to choice on each trial. The logistic regression model is given by:

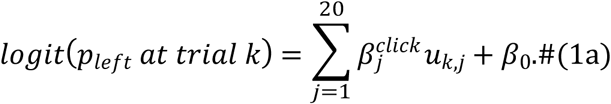

where *p*_*left*_ represents the probability of choosing left, *u*_*k*,*j*_ denotes *j*-th click side input (+ 1 for left click and ― 1 for right click) on trial *k*, {β^*click*^}^20^ is the integration kernel, and β_0_ is the overall decision bias.

### The rank-1 neural network model architecture and collective dynamics

We construct rank-1 network model with N neurons (N = 512 in this study) following the equation:

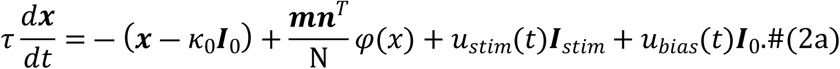

We generate the input embedding and recurrent connectivity matrices {*I*^(*i*)^,*I*^(*i*)^, *m*^(*i*)^,*n*^(*i*)^ of the rank-1 network by sampling neurons from a multi-variate gaussian i.i.d.:

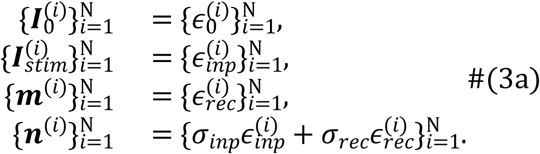

where ∈_0_,∈_*inp*_ and ∈_*rec*_ are independent white noises that account for connectivity in resting state, stimulus input and recurrent input respectively, with ∈_0_,∈_*inp*_,∈_*rec*_ ∼ *N*(0,*I*).

The hidden state of the network *x* explores a low-dimensional subspace spanned by the orthogonal column vectors *I*_0_,*I*_*stim*_,*m*:

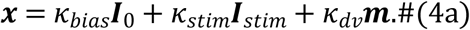

where *κ*_*bias*_,*κ*_*stim*_ and *κ*_*dv*_ are variables describing the collective dynamics of the network, i.e.,

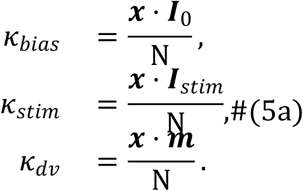

Together with (Eq. 2a), we derive the differential equations of collective variables as:

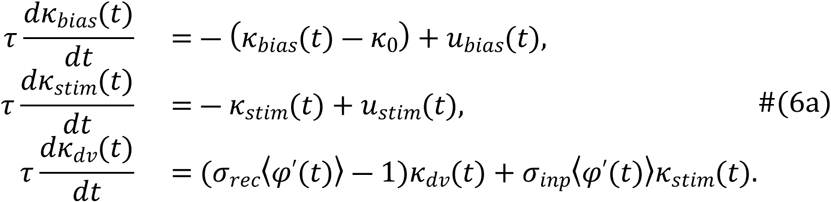

where ⟨φ′(*t*)⟩ represents the average single-neuron gain.

If N is sufficiently large, the average neuron gain can be approximated as:

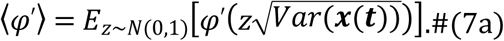

where *Var*(*x*(*t*)) is the variance of hidden state given by:

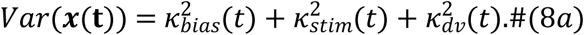

### Estimation of the integration kernel of rank-1 network models

By introducing *R*(*t*) = σ_*rec*_⟨φ^′^(*t*)⟩ ― 1 and S(*t*) = σ_*inp*_⟨φ^′^(*t*)⟩ as simplification, we can rewrite the collective dynamics of decision variable as the following equation:

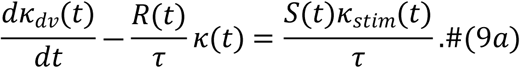

Multiply both side of the differential equation by the integration factor 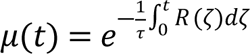 and simplifies the left-hand side of the equation as a total differential, we obtain:

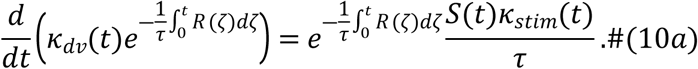

Integrating both sides from 0 to stimulus end *T*, and use the boundary condition *κ*_*dv*_(0) = 0, we have:

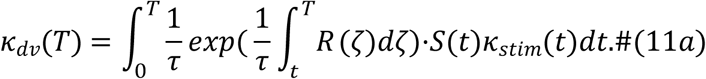

This provides the explicit form of integration kernel.

### Maximum likelihood estimate

To evaluate how well a participant or a model perform the Poisson-Click task, we use the maximum likelihood function. The likelihood for a set of participant data is given by:

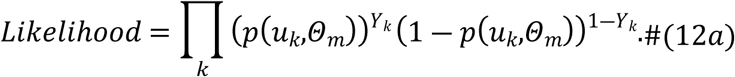

where {*u*_*k*_,Y_*k*_} represents the full set of participant’s data composed of click input time series *u*_*k*_ (+ 1 for left click and ― 1 for right click) and choices Y_*k*_(1 for left choice and 0 for right choice), θ_m_ denotes the model parameters. *p*(*u*_*k*_,θ_m_) is the probability of choosing left given click input time series *u*_*k*_and model parameters θ_m_.

The model assumes that decisions are made based on a properly scaled decision variable of the rank-1 network, with an overall decision bias β to either left or right side and an additional offset parameter μ for the integration kernel. The probability of choosing left at trial *k* is given by:

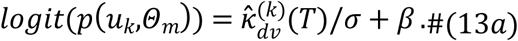

where 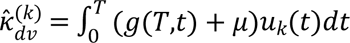

We choose model parameters that best fit participants’ choices by minimizing the negative loglikelihoods:

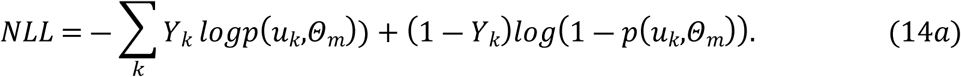

### Optimization of model parameters

After generating network models, we then optimize model parameters to fit human data. The parameters to be optimized include: {A,μ,β,*κ*_0_,*u*_0_,σ_*inp*_,σ_*rec*_}. The optimization process involves minimizing the negative log-likelihood of participants’ choices using the gradient descent algorithm. For each human participant, we start with 20 randomly sampled initial points in the parameter space. We then apply the gradient descent algorithm to adjust the parameters and minimize the negative log-likelihood. The optimization is conducted for a maximum of 2000 epochs to ensure convergence and accurate fitting of the model to the observed behavior.

Similarly for divisive normalization model and leaky competitive accumulators, we also optimize model parameters by minimizing negative log-likelihoods of participants’ choices. We use fmincon.m function from Matlab’s optimization toolbox with interior-point algorithm. Each human participant’s behavior is fitted by randomly sampling 360 starting points in the parameter space of each model respectively.

### Analysis of single neuron dynamics

To analyze the dynamics of single neurons in the rank-1 network models, we estimate how click input, choice, and time influence neural responses. This process involves fitting the activity of each neuron using logistic regression and interpreting the effects of collective variables on single neuron dynamics.

Here’s a detailed breakdown of the analysis: Firstly, we simulate the model with a dataset of 4096 artificial poisson-click trials. The activity r_*i,k*_(*t*) of neuron *i* on trial *k* is normalized to [0,1] with a linear mapping. The normalized neuron activity is denoted as r_*i,k*_(*t*). We model the normalized neuron activity r_*i,k*_(*t*) using logistic regression to account for the effects of time, choice, and stimulus input:

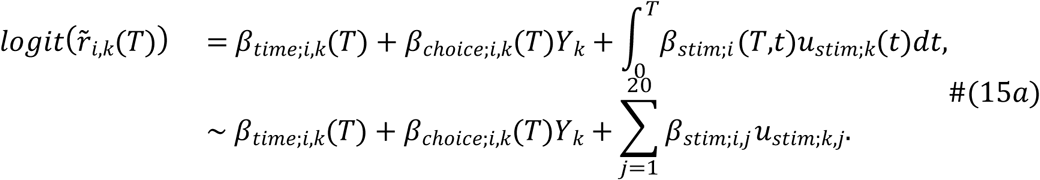

where Y_*k*_indicates participant’s choice on trial *k* (+ 1 for left choice and 0 for right choice). The regression coefficient β_time;*i,t*_(*T*), β_choice;*i,k*_(*T*) and β_*stim*;*i*_(*T,t*) represents the modulation of neuron *i* due to time, choice and stimulus input, respectively.

The activity of neuron *i* can be described as the combination of collective variables:

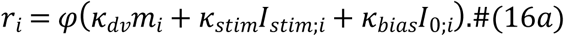

This indicates that the temporal weighting of the stimulus for each neuron consists of evidence accumulation and input filtering, each with distinct temporal characteristics.

The temporal weighting of collective variable *κ*_*dv*_(*T*) and *κ*_*stim*_(*T*) are:

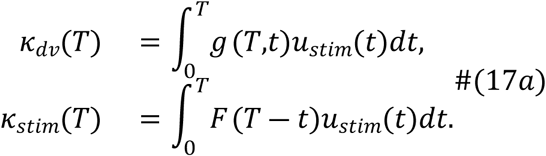

where *g*(*T,t*) and F(*T* ― *t*) are the integration and input kernels, respectively.

Using the integration kernel *g*(*T,t*) and input kernel F(*T* ― *t*), we fit the normalized neuron activity r_*i,k*_(*T*) with:

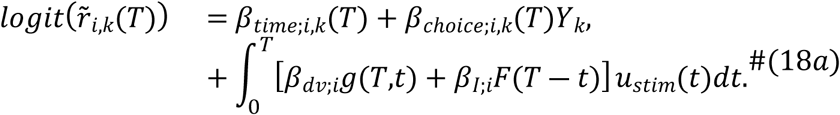

Here, β_*dv*;*i*_ and β_*I*;*i*_ account for the modulation of neuron *i* according to evidence accumulation and input signal processing.

The estimated single neuron integration kernel *g*_*i*_(*T,t*) is computed as:

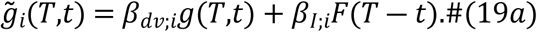

## Data and Code Availability

The data used in this study were obtained from Keung et al., as published in their article “Regulation of evidence accumulation by pupil-linked arousal processes” ([doi:10.1038/s41562-019-0551-4]). The dataset is publicly available from Open Science Framework (OSF) at https://osf.io/37yk8/. The code supporting the findings of this study will be made publicly available on Github upon publication of this article.

## Acknowledgment

We thank G. Okazawa for the critical comment on the manuscript. Funding: This study was supported by STI2030-Major Project 2021ZD0204105 (B.M.), National Science Foundation of China 32271149 (B.M.) and 12174007 & 12090054 (F.L.).

**Supplementary Notes.** The Supplementary Notes include 4 sections.

**S1 Fig.** Individual human behavior categorization. Black lines are individual integration kernel by regression of Eq. 1a. Colored lines are grouped kernel shapes with polynomial function fitting with Eq. 1s. Participants are grouped by kernel shapes colored with their indices marked on lower left.

**S2 Fig.** Rank-1 network model implements monotonic (a.) and non-monotonic integration kernels (b.) with flexible solutions. The left and right three columns are effective input strength, effective temporal factor and kernel estimation respectively. Black lines in the third column are kernels by regression of Eq. 1a. Red lines are estimated kernels by Eq. 11a.

**S3 Fig.** Theoretical prediction of integration kernels matches model simulation of rank-1 network with linear readout from neural firing rates. Black lines indicate integration kernel by regression of Eq. 1a. Red lines are estimated integration kernels by Eq. 3s.

**S4 Fig.** Individual integration kernels generated by best fitted rank-1 network model comparing to human behavior. Black and red lines are individual integration kernels by regression (Eq. 1a) over human participants’ choices and model simulation choices (Eq. 4), respectively.

**S5 Fig.** Individual psychometric curves generated by rank-1 network model comparing to human psychometric curves.

**S6 Fig.** Distribution of rank-1 network model parameters that best fit to human participants. The first row is boxplots of noise σ, overall decision bias β, offset for clicks μ. The second row is scatterplot of network connectivity parameters in pairs: {*K*0, ^*u*^0^,σ^*inp*^,σ^*rec*}.

**S7 Fig. Individual integration kernels generated by Leaky Competitive Accumulator compared to human integration kernels.** Best fitted parameters were obtained using fmincon.m function from Matlab’s optimization toolbox.

**S8 Fig.** Individual psychometric curves generated by Leaky Competitive Accumulator compared to human psychometric curves.

**S9 Fig.** Individual integration kernels generated by Divisive Normalization Model compared to human integration kernels. Best fitted parameters were obtained using fmincon.m function from Matlab’s optimization toolbox.

**S10 Fig.** Individual psychometric curves generated by Divisive Normalization Model compared to human psychometric curves.

**S11 Fig.** Neural integration kernels by regression over the normalized neural activity of the last time point compared to all time points.

**S12 Fig.** Neural integration kernel compared to mean-field theory approximation. Additional 24 neurons are marked on upper left scatter plot with red color.

**S13 Fig.** Single neuron integration kernels under *Softplus* activation. a. Single neuron kernel compared to reconstructed integration kernel with behavior and input kernel. b. Correlation between network connectivity weights and regression coefficients.

## Notes

### Competing Interest Statement

The authors have declared no competing interest.

